# XMAP215 and γ-tubulin additively promote microtubule nucleation in purified solutions

**DOI:** 10.1101/2020.05.21.109561

**Authors:** Brianna R. King, Michelle Moritz, Haein Kim, David A. Agard, Charles L. Asbury, Trisha N. Davis

## Abstract

Microtubule nucleation is spatiotemporally regulated in cells by several molecules, including the template γ-tubulin and the polymerase XMAP215. Recently, XMAP215 and the γ-tubulin ring complex were reported to function synergistically, and this synergy was hypothesized to be due to direct binding between XMAP215 and γ-tubulin. Here, we address this hypothesis by 1) probing domain requirements for XMAP215 to promote microtubule nucleation and 2) testing whether XMAP215 functions synergistically with γ-tubulin in the absence of the other ring complex proteins. We confirm that γ-tubulin and XMAP215 are classically defined nucleators that reduce the nucleation lag seen in bulk tubulin assembly. Then, using deletion constructs, we show that XMAP215’s ability to nucleate microtubules in purified solutions correlates with its ability to elongate existing microtubules and does not depend on the number of TOG domains. Finally, we show that XMAP215 and γ-tubulin promote αβ-tubulin assembly in an additive, not synergistic, manner. Thus, their modes of action during microtubule nucleation are distinct, and the synergy reported between XMAP215 and the γ-tubulin ring complex is not due to γ-tubulin alone.

## Introduction

Spontaneous nucleation of a new microtubule from nanometer-sized soluble αβ-tubulin dimers is an event that is currently impossible to observe directly. Tubulin dimers must associate longitudinally and laterally in order to form the final product: a hollow tube of side-by-side protofilaments. Models of the spontaneous microtubule nucleation pathway have disagreed on the number of rate-determining steps and key intermediates in the nucleation pathway [1]–[3].

More recently, studies of nucleation have centered on the γ-tubulin ring complex (γ-TuRC), which is essential for microtubule formation across organisms [4]–[9]. The γ-TuRC stably binds the microtubule minus end and is thought to act as a template onto which αβ-tubulin dimers associate [10]–[12], theoretically promoting lateral interactions between adjacent αβ-tubulin dimers [13]. However, increasingly higher resolution structures of purified γ-TuRC show that the template is not perfect—the 13-member γ-tubulin ring presented by the complex does not perfectly match the geometry of the microtubule [14]–[16]. Further, the fraction of purified γ-TuRC that nucleates *in vitro* is consistently low [15], [17]–[19]. It is hypothesized that, in order to nucleate well, γ-TuRC must be activated *in vivo* through interactions with other proteins [20]–[22].

There is a rapidly growing body of evidence that at least one of the crucial co-factors for the γ-TuRC is the XMAP215 family of microtubule polymerases. The XMAP215 family promotes microtubule growth from various templates, including the γ-TuRC, isolated centrosomes, and GMPCPP-stabilized seeds [15], [19], [23]–[26], and it also promotes microtubule nucleation in reactions without any template [23], [27]–[30]. Across these studies, the role of XMAP215 in nucleation was generally thought to be due to its polymerase activity. Recently, however, it was reported that XMAP215 and the γ-TuRC promote microtubule nucleation in a manner that is greater than the sum of their individual effects [15], [25], [26]. The mechanism underlying this reported synergy is unknown, although Thawani, et al. [25] hypothesized it is due to a direct binding interaction between the unstructured C-terminal tail of XMAP215 and γ-tubulin.

Here, we sought to test the hypothesis that direct interactions between XMAP215 and γ-tubulin, in the absence of other ring complex proteins, are sufficient for synergistic action. We use recombinantly expressed and purified protein, as well as the classical turbidity assay, to quantitatively examine these interactions in the most simplified system possible. We first show that γ-tubulin alone forms laterally-associated arrays that reduce the nucleation lag associated with αβ-tubulin assembly. We also confirm that XMAP215 alone reduces the nucleation lag and probe its mechanism of action using domain analysis, finding that nucleation activity strongly correlates with polymerase activity. We then explore the nucleation activity of the γ-tubulin arrays together with XMAP215. Across a range of concentrations of XMAP215 and across XMAP215 deletion constructs, we fail to find synergy with γ-tubulin. Rather, we find additive action under all conditions, indicating that the modes of action of XMAP215 and γ-tubulin during microtubule nucleation are independent, and that any synergistic behavior must arise from unique interactions made in the context of the γ-TuRC.

## Results and Discussion

### γ-tubulin and XMAP215 each promote microtubule nucleation in the classical turbidity assay

Nucleation of a new microtubule is impossible to observe directly. Light microscopy can visualize dynamics of individual microtubule formation events, but it is complicated by the requirement of fluorophore-labeled αβ-tubulin heterodimers. In order for αβ-tubulin to assemble, only a small fraction of αβ-tubulin may carry a fluorophore; this not only makes early nucleation intermediates impossible to visualize, but it also indicates that unlabeled and labeled αβ-tubulin assemble differently.

Light scattering, on the other hand, uses unlabeled αβ-tubulin heterodimers to monitor formation of small nucleation intermediates. This assay, performed in bulk, is classically used in the study of polymerization dynamics [31], [32]. The initial delay or lag between the start of the reaction and when the turbidity reaches one-tenth of its maximum provides a sensitive measure of polymer nucleation efficiency [1], [3].

To establish the light scattering assay in our hands, we first tested an accepted αβ-tubulin nucleator, γ-tubulin. At higher concentrations (∼250 nM and above), purified γ-tubulin oligomerizes, forming laterally-associated arrays with plus-ends of γ-tubulin oriented outward, as revealed by negative stain electron microscopy (Figure 1, A and B), and as seen previously in γ-tubulin crystals [13]. We first confirmed that these arrays alone do not scatter light and that they promoted formation of microtubules rather than other non-canonical tubulin polymers (Supplementary Figure 1). We monitored the effect of γ-tubulin arrays on αβ-tubulin assembly, finding that they strongly decreased the nucleation lag compared to αβ-tubulin alone (Figure 1C). In contrast, lower concentrations of γ-tubulin that failed to form laterally-associated arrays only weakly promoted nucleation (Figure 1C). To quantify the nucleation activity of γ-tubulin for our comparisons with XMAP215, we used 300 nM γ-tubulin, a sufficiently high concentration for array formation. At this concentration, the γ-tubulin arrays decreased the time to reach 10% polymer formation, hereafter known as the nucleation lag, by approximately 50% (Figure 2A).

**Figure 1.**
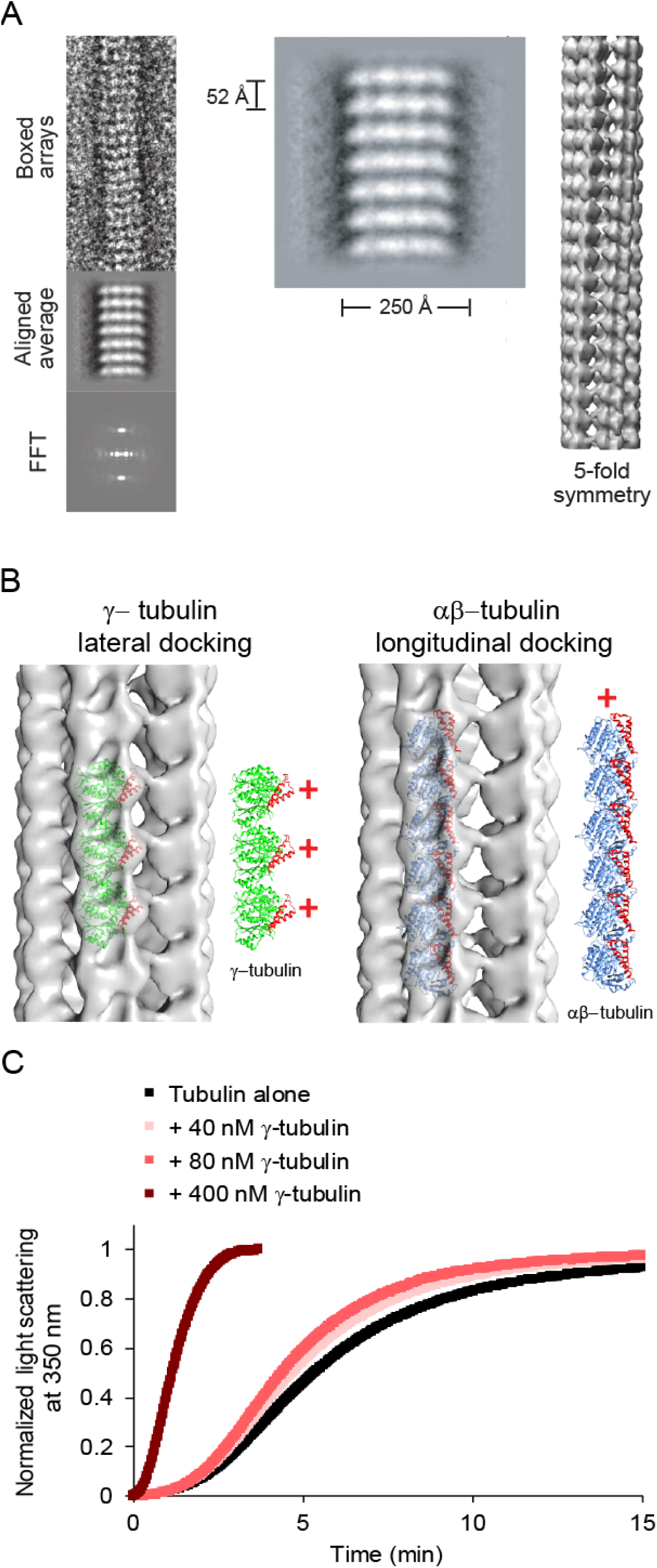
Negative-stain electron microscopy reveals that γ-tubulin forms arrays that are distinct from microtubules. **(A)** γ-tubulin arrays were boxed and averaged for helical reconstruction. 2D averages of the arrays showed a ∼52 Å repeat unit and ∼250 Å width. Helical reconstruction reveals 5-fold symmetry in the array, with a hollow center. **(B)** Left: Docking of the human γ-tubulin crystal structure (1Z5W) into the 3D reconstruction revealed that γ-tubulins are laterally associated along the long axis of the arrays. The lateral packing of γ-tubulins is similar to that observed in crystals [13]. γ-Tubulins in the arrays are oriented with their plus-ends outward (red +). Helices 3 and 4 are red in the models to illustrate the orientation of the plus-end of the γ- or αβ-tubulins. Right: Docking of αβ-tubulins (1JFF) as a proxy for a hypothetical longitudinal assembly of γ-tubulins. The periodicity of the longitudinal model does not match that of the EM map. **(C)** Characterization of 40, 80, or 400 nM γ-tubulin in the turbidity assay with 12 μM αβ-tubulin.

**Figure 2.**
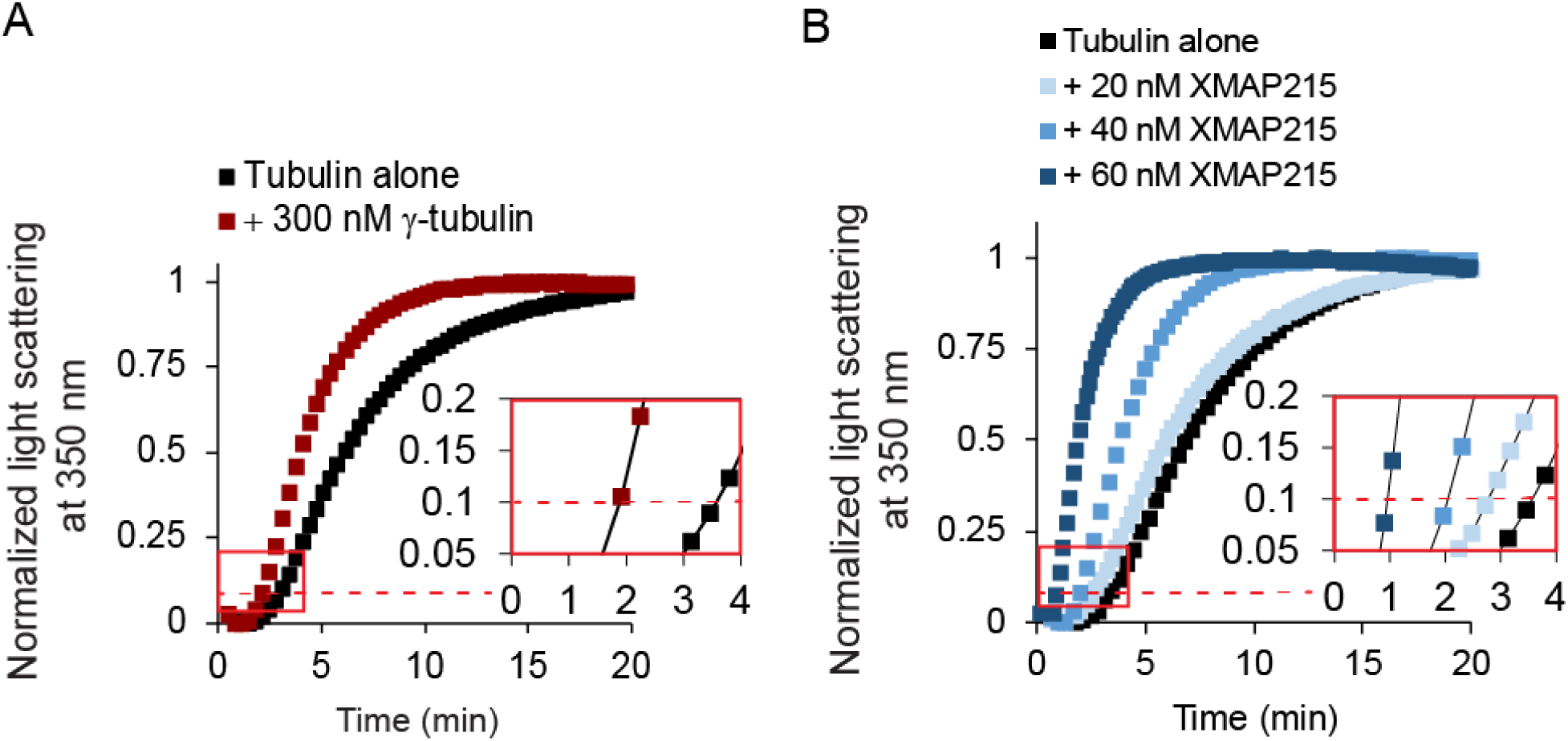
Laterally-associated γ-tubulin arrays and XMAP215 are classically-defined nucleators. (A) Representative turbidity assay at 15 μM free αβ-tubulin, with γ-tubulin added at 300 nM. Red dashed line represents 10% of the maximum light scattering. Inset shows same data from red box plotted on larger scale for clarity. (B) Representative tubulin assembly assay for 20 nM, 40 nM, or 60 nM XMAP215 with 15 μM free tubulin. XMAP215 promotes spontaneous tubulin assembly in a concentration-dependent manner. Red dashed line represents 10% of the normalized maximum light scattering. Inset shows same data from red box plotted on larger scale for clarity.

We then tested the nucleation activity of XMAP215, finding significant effects. The nucleation lag decreased monotonically with increasing concentrations of XMAP215, with a decrease of 75% at the highest concentration (60 nM, Figure 2B). This provided independent confirmation that XMAP215 is a nucleator, with similar potency to γ-tubulin arrays.

### XMAP215 functions as a microtubule polymerase during microtubule nucleation

XMAP215 has five TOG domains as well as a basic region referred to as the microtubule binding domain (MTbd). Previous domain analyses of XMAP215 have shown that two separate types of domains are required for polymerase activity: 1) TOGs 1 and 2, which bind free αβ-tubulin dimers with high affinity, and 2) a microtubule-lattice binding region, which can be provided by either the MTbd or by TOGs 3 and 4 [26], [30], [33], [34]. To test if the XMAP215 domain requirements for nucleation activity are the same as those for polymerase activity, we created an array of constructs with varying polymerase activities. In our constructs, we maintained the presence of TOGs 1 and 2, which are essential for polymerase activity, and varied the domains required for localization to the microtubule lattice (Figure 3A).

**Figure 3.**
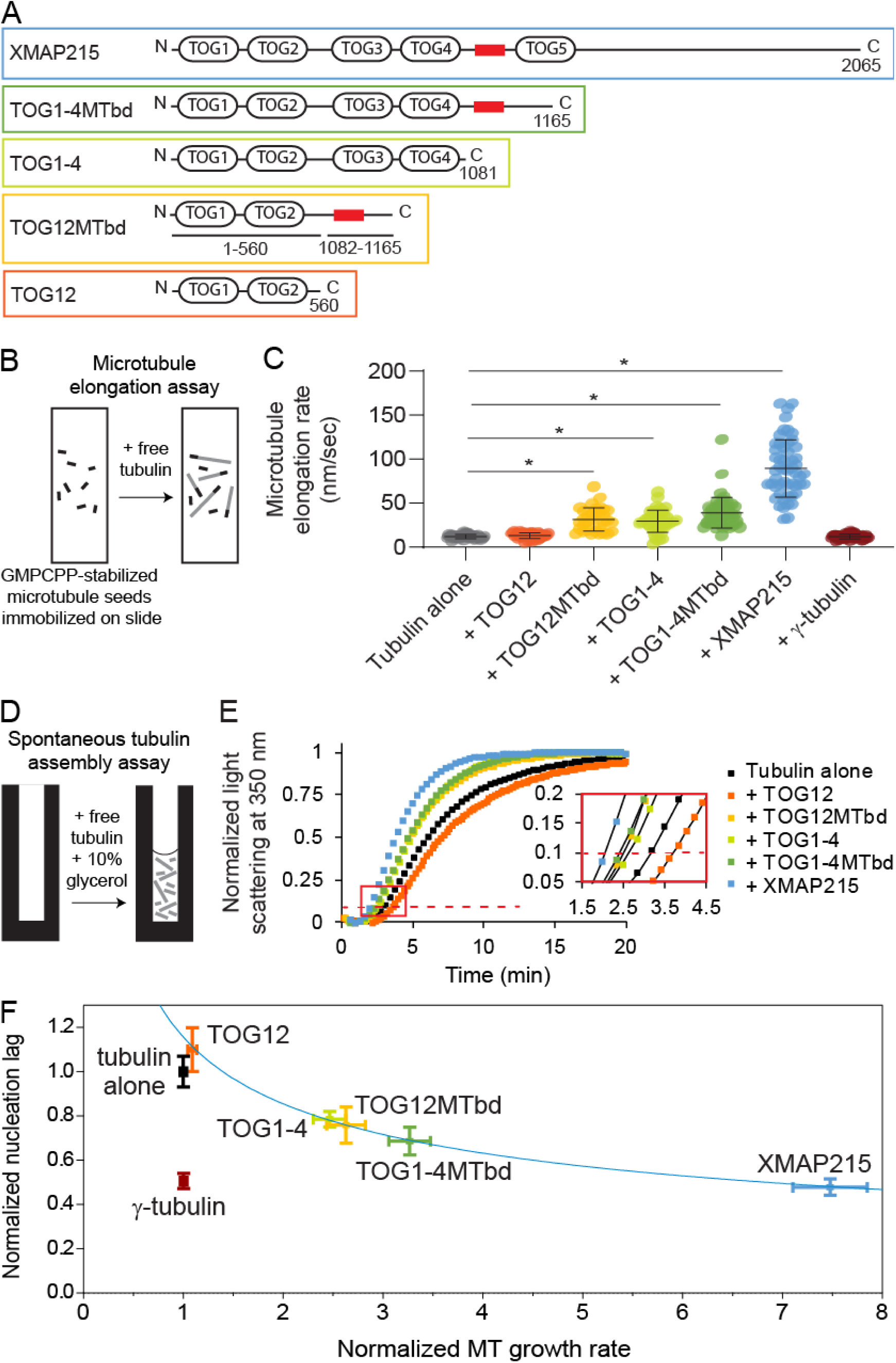
The microtubule nucleation activity of XMAP215 correlates strongly with its polymerase activity. **(A)** XMAP215 and XMAP215 deletion constructs used in this work. The red box represents the basic MTbd. **(B)** Schematic of the assay used to measure tip growth rates from immobilized GMPCPP-stabilized microtubule seeds by microscopy. **(C)** Microtubule elongation rates at 12 μM free αβ-tubulin, with full-length XMAP215 or the indicated deletion constructs added at 40 nM, or with γ-tubulin added at 300 nM. Error bars represent standard deviation (N = 41, 35, 33, 41, 49, 53, 41 for each respective condition) microtubules measured for each condition). Asterisk represents p<0.0001, two-sided t-test. **(D)** Schematic of the assay used to measure non-templated tubulin assembly by turbidity. **(E)** Representative turbidity assay at 15 μM free αβ-tubulin, with XMAP215 or the indicated deletion constructs added at 40 nM. Red dashed line represents 10% of the maximum light scattering. Inset shows the data indicated by the red box re-plotted on a larger scale for clarity. **(F)** Nucleation activity plotted against polymerase activity. Data for XMAP215 and its deletion constructs, each added at 40 nM, are fit by a power curve with the equation f(x) = 1.16x^-0.436^ (see Methods). Normalized microtubule growth rate means ± SEM represent the data from Figure 3C normalized to the growth rate of tubulin alone. Relative nucleation activity for each construct is represented by delay between the start of the polymerization reaction and when the turbidity reached 10% of its maximum, divided by the delay measured in controls with αβ-tubulin alone. Normalized nucleation rate means ± SEM represent the continuous turbidity reaction data of experimental conditions normalized to the nucleation lag of tubulin alone (N ≥ 12, 7, 6, 4, 6, 5, and 6 for tubulin alone, γ-tubulin, TOG12, TOG12MTbd, TOG1-4, TOG1-4MTbd, and XMAP215, respectively).

To examine the polymerase activities of the XMAP215 constructs, we measured microtubule elongation rates for individual microtubule plus-end tips assembling continuously from GMPCPP-stabilized seeds (Figure 3, B and C). As seen in previous studies, addition of full-length XMAP215 strongly accelerated microtubule elongation, increasing the plus-end growth rate by up to 10-fold over αβ-tubulin alone. Addition of TOG1-4MTbd, TOG12MTbd, or TOG1-4 at identical concentrations also accelerated growth, but increased the rate more modestly, by 2.5- to 3.3-fold. Addition of TOG12 had no significant effect. The necessity of TOGs 1 and 2 plus either the MTbd or TOGs 3 and 4 for polymerase activity confirms previous findings [26], [33], [34].

We then tested the activity of our XMAP215 constructs in microtubule nucleation. Three of the deletion constructs added at identical concentrations had moderate nucleation activity compared to full-length XMAP215: TOG1-4MTbd, TOG12MTbd, and TOG1-4. Of these, TOG1-4MTbd had the largest effect, reducing the nucleation lag by 31% compared to αβ-tubulin alone (Figure 3E). TOG12MTbd and TOG1-4 promoted nucleation similarly; each reduced the nucleation lag by ∼25%. Only one of the deletion constructs we tested, TOG12, showed no effect on nucleation. Notably, the activity of the constructs in nucleation was not dependent on the number of TOG domains.

Similarities in the domain requirements for nucleation versus polymerase activity were striking. Deletion of TOG5 and the C-terminus of XMAP215 significantly reduced both polymerase and nucleation activity. Further reduction of microtubule lattice affinity, by deleting either TOGs 3 and 4 (TOG12MTbd construct) or the basic region following TOG4 (TOG1-4 construct), predictably reduced polymerization activity and likewise reduced nucleation activity. Elimination of microtubule lattice affinity altogether (TOG12) eliminated both activities. Plotting the turbidity and tip growth data against one another illustrates that the results from these two experiments are correlated (Figure 3F). Their relationship suggests that XMAP215’s role during nucleation is strictly as a microtubule polymerase. That is, the nucleation activity of XMAP215 does not depend on any particular lattice-binding domain or on the number of TOG domains, but rather it depends on microtubule polymerase activity. Polymerase activity and nucleation activity both require only TOGs1 and 2 and a lattice binding region.

### γ-tubulin promotes microtubule nucleation without affecting elongation rate

For comparison with XMAP215, it was instructive to also compare the effects of γ-tubulin in the tip growth and turbidity assays. Addition of purified γ-tubulin alone had no effect on plus-end growth rates (Figure 3C), which is not surprising since γ-tubulin caps minus ends and does not associate with microtubule plus ends. However, the same concentration of γ-tubulin dramatically reduced the nucleation lag in the turbidity assay, by 50% (Figures 2A and 3F). These observations confirm that γ-tubulin promotes microtubule nucleation without affecting microtubule plus-end elongation, consistent with its expected role as a minus-end-binding template.

### XMAP215 and γ-tubulin function additively during microtubule nucleation

If XMAP215 functions strictly to elongate nucleation intermediates during microtubule formation, then its role should not overlap with γ-tubulin, and thus the effects of XMAP215 and γ-tubulin arrays in the turbidity assay should be additive. In contrast, if XMAP215 and γ-tubulin arrays function synergistically (or redundantly), they should produce a larger (or smaller) effect when combined than the sum of their independent contributions.

To distinguish the type of relationship XMAP215 and γ-tubulin arrays display, we calculated the effect on nucleation lag that would occur in a purely additive scenario (Supplementary Figure 2 shows example calculations). Then, over the previously tested concentrations of XMAP215, we measured the combined effects of XMAP215 and γ-tubulin arrays (Figure 4A). We compared the observed nucleation lags with the predicted nucleation lags, and found they closely matched across all XMAP215 concentrations (Figure 4B).

**Figure 4.**
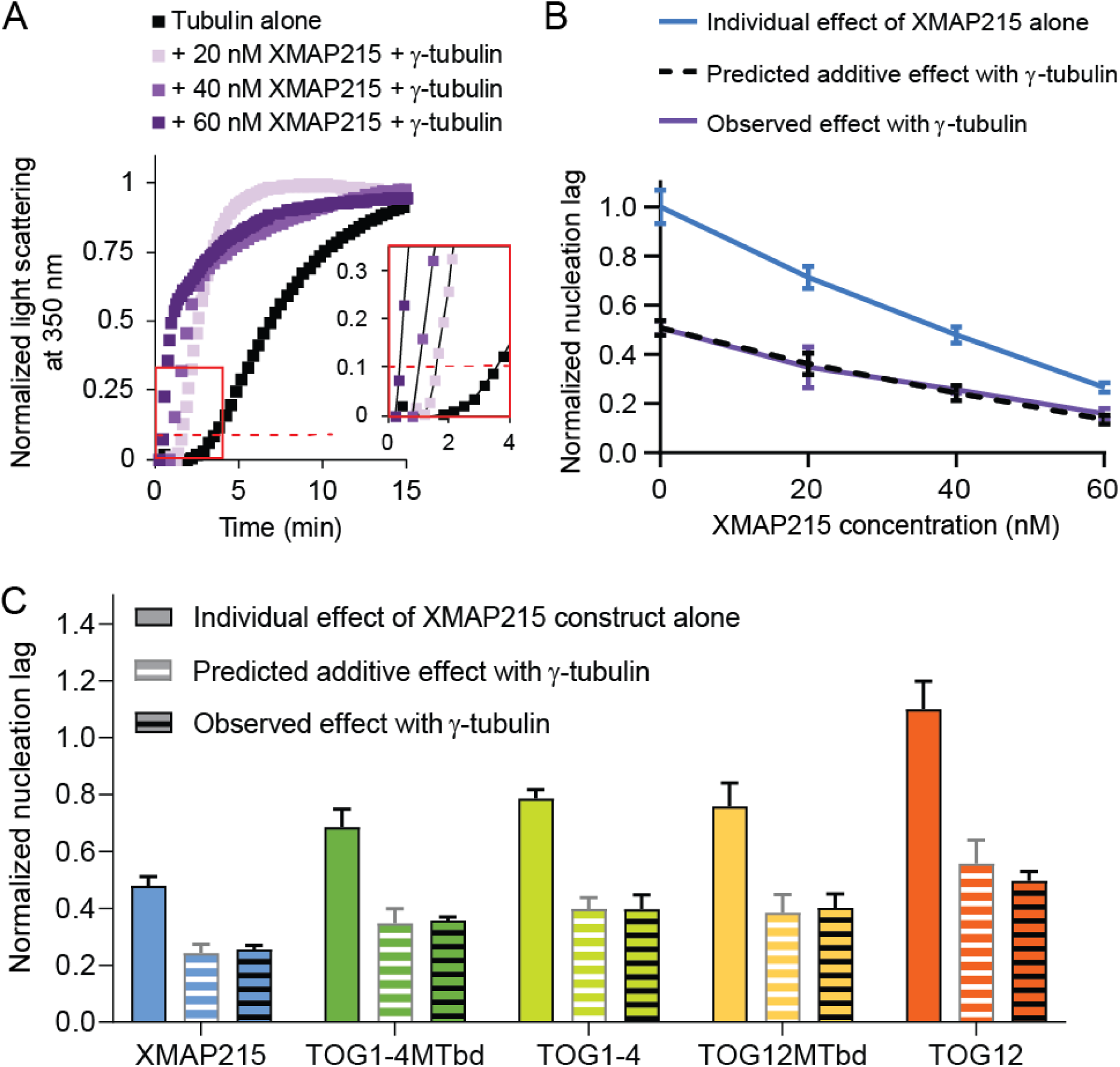
XMAP215 and γ-tubulin promote nucleation additively in pure tubulin solutions. **(A)** Representative tubulin assembly assay for 20 nM, 40 nM, or 60 nM XMAP215 in combination with 300 nM γ-tubulin with 15 μM free tubulin. Red dashed line represents 10% of the normalized maximum light scattering. Inset shows same data from red box plotted on larger scale for clarity. For 40 nM and 60 nM XMAP215 in combination with 300 nM γ-tubulin, microtubule bundling occurs at later time points, which causes a difference in the shape of the curve of tubulin assembly. See Supplementary Figure 3. **(B)** XMAP215 functions additively with γ-tubulin at all concentrations measured (see Supplementary Figure 2). Data plotted are mean values ± SEM (N = 12, 4, 6, 4 for 0 nM, 20 nM, 40 nM, and 60 nM XMAP215, respectively; N = 7, 4, 4, and 4 for 0 nM, 20 nM, 40 nM, and 60 nM XMAP215 + γ-tubulin, respectively). **(C)** XMAP215 deletion constructs all show additive action with γ-tubulin (see Supplementary Figure 2). Data plotted for each XMAP215 deletion construct are mean values ± SEM replotted from Figure 3F. Data plotted for each XMAP215 deletion construct + γ−tubulin are mean values ± SEM (N = 4 for each condition).

To further test if XMAP215 and γ-tubulin act in an additive manner, we measured the combined effect of each of our XMAP215 deletion constructs with γ-tubulin. In every case, the nucleation lag observed when the two were combined was not significantly different from the predicted additive value (Figure 4C). Thus, the removal of various domains from XMAP215 does not allow for either synergy or redundancy with γ-tubulin. Overall, the close correspondence of the measured nucleation lags with the predicted additive values, across different XMAP215 concentrations and across the different deletion constructs, shows that XMAP215 and γ-tubulin function additively in the absence of other ring complex proteins.

During polymerization, XMAP215 increases the on-rate for tubulin dimers associating with the microtubule tip [35]. Our results suggest that this is its primary contribution during microtubule nucleation as well. XMAP215’s effect on microtubule nucleation correlates with its polymerase activity, independent of the domains present. This provides support for a model of independent action by XMAP215 during microtubule nucleation, as suggested previously [19], [27], [36].

We show here that XMAP215 functions additively with laterally-associated arrays of γ-tubulin. That a microtubule polymerase, which promotes elongation, and a template, which promotes lateral interactions, act independently during the early stages of microtubule formation supports early nucleation models that include several rate-limiting steps [1], [3] and a more recent model that proposes microtubules form by progressively faster tubulin accretion [37]. Our results suggest that there must be at least two distinct types of rate-limiting transitions in the nucleation pathway. Given the theoretical activities of the template and the polymerase, we propose that two of these steps are (i) the formation of initial lateral αβ-tubulin bonds, which is specifically accelerated by γ-tubulin, and (ii) the formation of subsequent αβ-tubulin bonds, which is specifically accelerated by XMAP215.

Our results also show that γ-tubulin and XMAP215 do not show the synergistic interaction that was previously reported for the full γ-TuRC and XMAP215. Several possibilities could explain this difference. It is thought that the γ-TuRC is activated through a conformational change, and the reported interaction between the unstructured C-terminal tail of XMAP215 and γ-tubulin might be responsible for this. It is also possible that the full γ-TuRC offers higher affinity binding sites for this interaction with XMAP215 than arrays of γ-tubulin alone. Extra molecules of XMAP215 would increase the local concentration of αβ-tubulin near the template and thereby increase the rate of addition of αβ-tubulin. Alternatively, the arrangement of a γ-tubulin template into a ring might activate it for synergistic action by forming better substrates for XMAP215. Laterally-associated γ-tubulin arrays and γ-TuRCs both provide templates that should promote lateral interactions between αβ-tubulin dimers. However, the templates are not identical and each might promote a distinct ensemble of nucleation intermediates. According to this model, XMAP215 would act preferentially on nucleation intermediates that form on the 13-member γ-tubulin template provided by the ring complex.

## Methods

### Purification of tubulin

αβ-tubulin was purified from fresh bovine brain using a high-concentration PIPES buffer as previously described [38].

### γ-tubulin and XMAP215 expression plasmids

A human γ-tubulin construct with a C-terminal myc-His_6_ tag was previously described [39]. A construct with XMAP215 with a C-terminal GFP-His_7_ tag was previously described [40]. The fragment containing GFP was removed using SalI digestion (pRK009) and QuikChange was performed to produce TOG1-4MTbd (pRK087) and TOG1-4 (pRK031). pPW263, a bacterial expression plasmid containing TOG12 and three K loops with a C-terminal myc-His_6_ tag was previously described [33]. QuikChange was performed to produce TOG12MTbd (pRK074) and TOG12 (pRK056).

### Purification of γ-tubulin and XMAP215 constructs

γ-tubulin, XMAP215, TOG1-4MTbd, and TOG1-4 were expressed in Sf9 cells using the Bac-to-Bac system from Invitrogen (Thermo Fisher Scientific, Waltham, Massachusetts). Sf9 cells at a density 1-2×10^6^ cells/ml were infected with baculovirus. Cell pellets were harvested 48-72 hrs. post-infection. Infections were done both in-house, at the National Cell Culture Center (Minneapolis, MN), or at the Tissue Culture Core Facility, Univ. of Colorado Cancer Center, UCHSC at Fitzsimons (Aurora, CO). TOG12MTbd and TOG12 were expressed in *E. coli* Rosetta 2 cells with plasmid pRARE. All XMAP215 and TOG constructs were purified as previously described [41], and γ-tubulin was purified as previously described [13].

### Spontaneous tubulin assembly assay

The spontaneous assembly of purified bovine brain tubulin was measured by light scattering at 350 nm as described previously [1]. Tubulin was thawed on ice slurry (0°C) and pelleted in a TLA100 rotor at 90,000 rpm for 10 minutes to remove microtubule polymers and/or inactive tubulin. The supernatant was removed, kept at 0°C, and used for the assembly reactions. Experimental reactions were prepared in 1.7 mL Eppendorf tubes at 0°C. Immediately before measuring light scattering, tubulin was added at either 12 μM or 15 μM to experimental reactions at 0°C for a total volume of 150 µL, and then the entire reaction was moved to a pre-warmed cuvettes at 37°C in a DU 800 spectrophotometer (Beckman Coulter Life Sciences, Indianapolis, Indiana). The A_350 nm_ was recorded every 6-20 seconds on 2-6 samples at a time for greater than 40 minutes. Recording on the spectrophotometer was started prior to tubulin being added to experimental reaction mix. For XMAP215 and γ-tubulin, at least 7 experimental samples per condition were tested, for two independent purifications of each protein. For XMAP215 deletion constructs, at least 4 experimental samples were tested. All conditions were randomized by experiment across multiple days. Scaling analysis was performed by hand.

### Microtubule growth rate assay

Small flow chambers were constructed by adhering a KOH etched coverslip onto a glass slide with two strips of double-sided tape. Flow chambers were incubated with pre-warmed 1 mg/ml biotin-BSA (Vector Laboratories, Burlingame, California) for 15 minutes at room temperature, then washed with pre-warmed BRB80 (80 mM PIPES (EMD Chemicals Inc., Gibbstown, New Jersey), 120 mM KCl (Hach Company, Loveland, Colorado), 1 mM MgCl_2_ (Millipore Sigma, St. Louis, Missouri) and 1 mM EGTA (Millipore Sigma), pH 6.8). GMPCPP-stabilized microtubule seeds assembled from a mixture of biotinylated porcine tubulin, 7 or 14% of total, (Cytoskeleton, Inc., Denver, Colorado) and purified bovine tubulin were secured to the glass coverslip surface using 0.25 mg/mL avidin (Vector Laboratories) [42]–[45]. After 5 minutes, microtubule seeds were washed with BRB80 containing 1 mM GTP (Millipore Sigma), 8 mg/mL BSA (Millipore Sigma), and 1 mg/mL K-casein (Millipore Sigma), until reaction mixture was added.

Dynamic microtubule extensions were grown in the absence, or presence, of 40 nM XMAP215 constructs. Reaction mixture consisted of freshly thawed 12 µM purified bovine brain tubulin in microtubule growth buffer (BRB80, 1 mM GTP (Millipore Sigma), 5 mM DTT (Millipore Sigma), 25 mM glucose (Millipore Sigma), 200 µg/mL glucose oxidase (Millipore Sigma), 35 µg/mL catalase (Millipore Sigma)). Microtubule extensions were imaged by recording video-enhanced differential interference contrast (VE-DIC), which allowed us to visualize the dynamic microtubule tips. A field of view (FOV) was imaged for 15-20 minutes. 4-8 FOVs per flow chamber were analyzed per experiment.

Growing microtubule tips were manually tracked using MTrackJ. X-Y coordinates of the microtubule tips were projected onto a line corresponding to the microtubule long-axis. Projected tip positions were then plotted against time, and during episodes of steady growth were fitted to a line. The slopes of these line-fits were recorded as the growth velocities.

### Nucleation lag and elongation rate correlational analysis

Data was plotted and fit using Igor Pro software (WaveMetrics, Portland, Oregon). A power curve was chosen to fit the data, in consideration of the theoretical relationship between microtubule elongation and nucleation. As microtubule growth rate approaches zero, the nucleation lag should increase and approach infinity. In contrast, as microtubule growth rate approaches infinity, the nucleation lag should decrease and approach zero. In accordance with this last point, the horizontal asymptote was constrained to zero. Additionally, the curve was weighted in relation to the SEM for the normalized nucleation lag data.

### Electron microscopy and helical reconstructions

γ-tubulin arrays were prepared by diluting pure γ-tubulin to 0.5-1 μM out of 0.5 M KCl-containing buffer to ≤ 100 mM KCl, no GTP. Notably, arrays form at 4°C and 37°C, with or without GTP. For negative stained samples, carbon-coated grids were glow-discharged or plasma-cleaned prior to use, followed by application of 3-5 μl of protein solution for 60 secs. Grids were then washed in ddH_2_O and stained with filtered 1% uranyl acetate for 30 secs. Images were obtained on a Tecnai20 transmission electron microscope (FEI, Hillsboro, Oregon) at 62,000 magnification and 200kV in low-dose mode with the use of either a 1024×1024 pixel CCD camera or a 4096×4096 pixel CCD camera (Gatan, Pleasanton, California) at -1.3-1.5 μm defocus. For γ-tubulin array reconstructions, individual arrays 300 pixels wide were extracted from the original micrographs using EMAN2 Helixboxer [46], after rotation to vertical along the long array axis. SPIDER [47], [48] was then used to window the arrays along the long axis, shifting by 270 pixels for 90% overlap between adjacent images. The image stacks were then padded by 90 pixels and binned 2x for a final square image size of 195 pixels. Image stacks corresponding to individually picked arrays were aligned in 2D to initially characterize array symmetry. Using a simple rectangular initial reference image based on an unaligned average of the image stacks, 10 rounds of reference-based alignment and averaging were run to calculate a final average image. Three similar, long and well-ordered arrays consisting of 560 overlapping segments were chosen as an initial dataset for 3D reconstruction. A repeat distance of ∼ 52 Å was measured from the 2D averages and used as a starting parameter for 3D reconstruction. The IHRSR program was used as the 3D helical alignment algorithm [49]. 50 rounds of refinement were run for each experiment using a simple cylinder with a diameter similar to the width of the 2D averages of the images. Multiple runs of IHRSR were performed using different initial azimuthal angles. Several different point group symmetries were also applied. The best results were obtained using a 5-fold point group symmetry with a final refined azimuthal angle of 6.5° and a rise of 52.9 Å. The final volume was contoured to an estimated 8.4 MDa and x-ray structures of γ-tubulin trimers [13] were manually fit to the corresponding density.

### Microtubule pelleting assay

Experimental reactions were prepared as described in the spontaneous tubulin assembly, and incubated at 37°C for 30 min. The reactions were fixed by 1:4 dilution in 2% glutaraldehyde (Electron Microscopy Services, Hatfield, Pennsylvania), and 2 min incubation at room temperature. Dilution of samples (1:1200) before pelleting was required to differentiate individual microtubules. Pelleting onto coverslips was performed as previously described [50].

To visualize microtubules on coverslip, immunostaining was performed using anti-α-tubulin—FITC antibody (Millipore Sigma, St. Louis, Missouri). The slides were imaged using an AxioObserver Z1 microscope (Zeiss, Jena, Germany) equipped with an ORCA-Flash4.0 camera (Hamamatsu Photonics K.K., Bridgewater, New Jersey) and an α Plan-Apochromat 63X objective (1.46 NA) (Zeiss). Immunostained microtubules were imaged with 0.4 s exposures and were not binned with a final resolution of 2048 × 2048.

## Acknowledgments

We thank Albion Baucom and Koji Yonekura for help with reconstructions of the γ-tubulin arrays. We thank Per Widlund for XMAP215 expression plasmids and helpful discussion. This study was supported by grants from the National Institutes of Health grant P01 GM105537 to Mark Winey, Trisha Davis, David Agard, Charles Asbury, R35 GM130293 to Trisha Davis, R35 GM134842 to Charles Asbury, R35 GM118099 to David Agard, and T32 GM007270 to Brianna Rose King.

## Author contributions

BRK designed and performed research, analyzed the data, and wrote the manuscript. MM performed research, contributed experimental design, contributed materials, and edited the manuscript. HK performed research. DAA contributed experimental design, supervised research, and edited the manuscript. CLA contributed experimental design, supervised research, and edited the manuscript. TND contributed experimental design, analyzed data, supervised the research, and helped write the manuscript.

---

**Table 1.** Plasmids used in this study.

| Plasmid name | Protein expressed | Vector | References |
| --- | --- | --- | --- |
| γ-tubulin-myc-His | γ-tubulin-myc-His<sub>6</sub> | pFastBac | [39] |
| XMAP215-GFP-His | XMAP215-GFP-His<sub>7</sub> | pFastBac | [41] |
| pRK009 | XMAP215-His<sub>7</sub> | pFastBac | This study |
| pRK087 | TOG1-4MTbd-His<sub>7</sub> | pFastBac | This study |
| pRK031 | TOG1-4-His<sub>7</sub> | pFastBac | This study |
| pPW263 | TOG12-3Kloop-myc-His<sub>6</sub> | pTrcHis A | [33] |
| pRK074 | TOG12MTbd-His<sub>6</sub> | pTrcHis A | This study |
| pRK056 | TOG12-His<sub>6</sub> | pTrcHis A | This study |

