## Supplemental Figures for "XMAP215 and γ-tubulin additively promote microtubule nucleation in purified solutions"

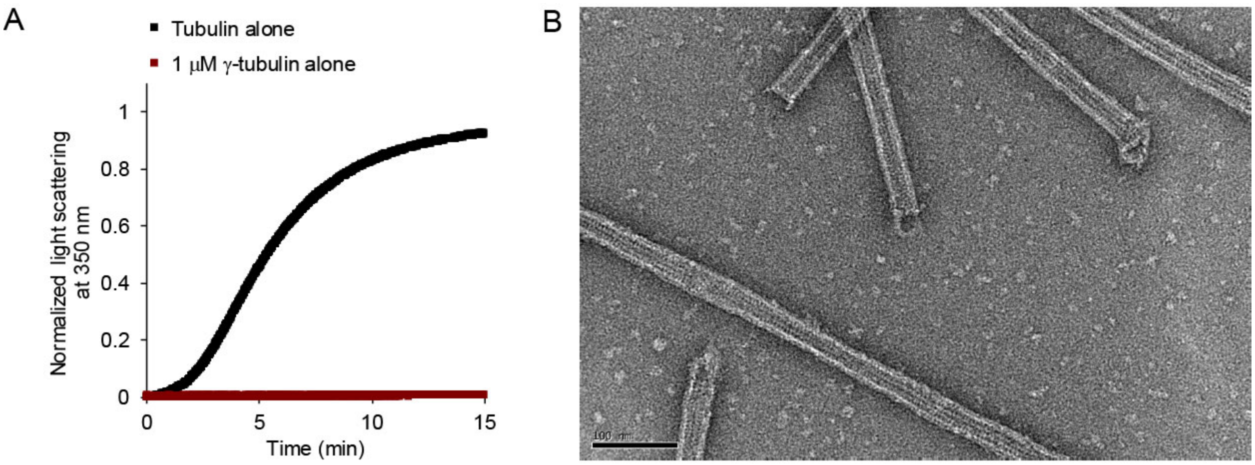


**Supplementary Figure 1.** γ-tubulin arrays promote microtubule formation in a light scattering assay.

**(A)** Turbidity assay at 12 μM free αβ-tubulin or 1 μM γ-tubulin, showing that γ-tubulin arrays alone do not scatter light under our experimental conditions.

**(B)** Negative-stain electron microscopy image of a sample from a light scattering assay using 1 μM γ-tubulin and 5 μM αβ-tubulin, showing that γ-tubulin arrays support the formation of microtubules, as opposed to other types of tubulin polymers.

| Condition | Normalized nucleation lag |
|---|---|
| Tubulin alone | 1 |
| + 300 nM γ-tubulin | 0.507 |
| + 20 nM XMAP215 | 0.749 |

Normalized nucleation lag with 300 nM γ-tubulin alone: 0.507

x

Normalized nucleation lag with 20 nM XMAP215 alone: 0.749

Predicted additive effect: 0.380

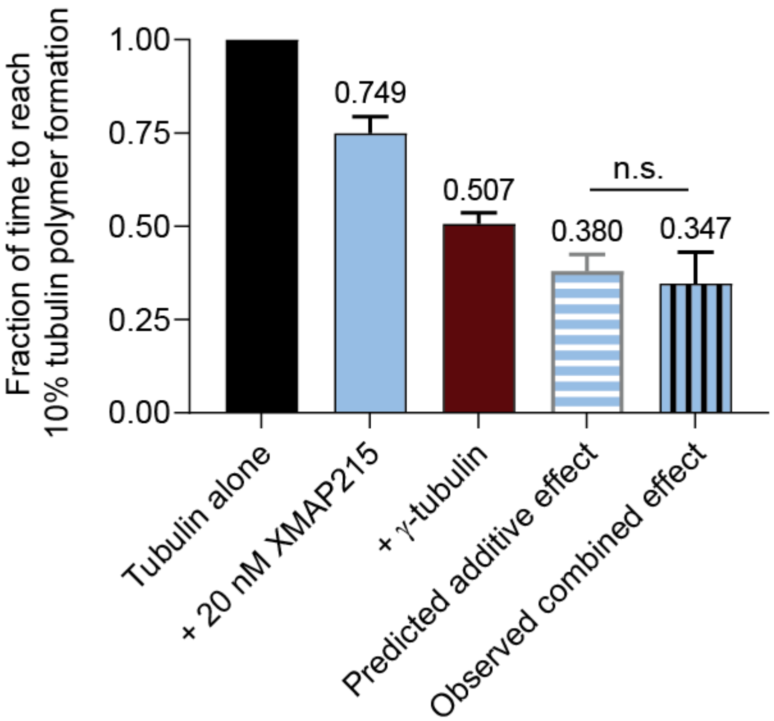


**Supplementary Figure 2.** Example calculation for predicted additive effect.

Data plotted are mean values ± SEM (N = 12 for tubulin alone, N = 4 for all other conditions). Significance was tested, two-sided t-test.

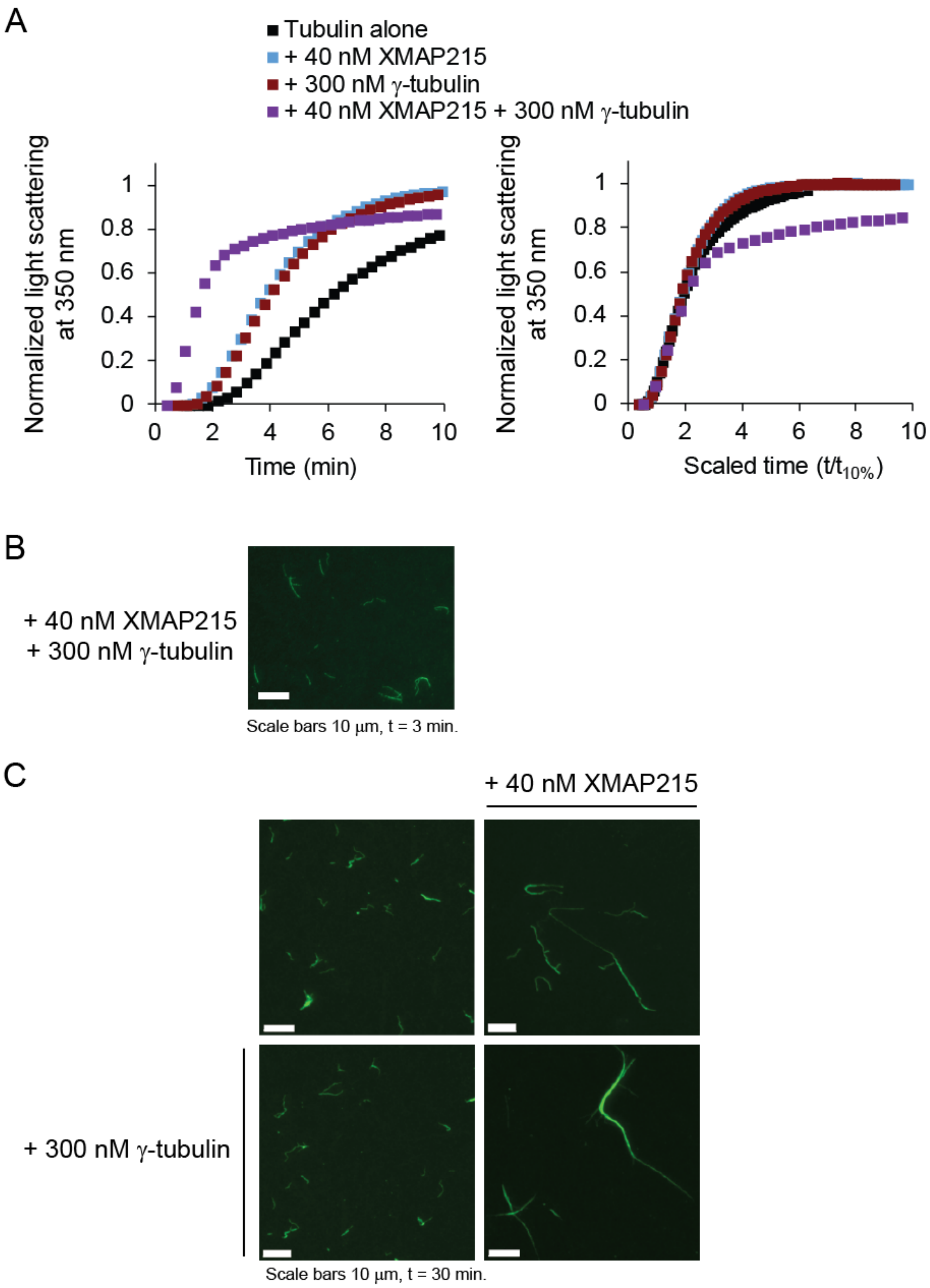
A
Tubulin alone
+ 40 nM XMAP215
+ 300 nM γ-tubulin
+ 40 nM XMAP215 + 300 nM γ-tubulin
Normalized light scattering at 350 nm
Time (min)
Scaled time (t/t10%)
B
+ 40 nM XMAP215
+ 300 nM γ-tubulin
Scale bars 10 µm, t = 3 min.
C
+ 40 nM XMAP215
+ 300 nM γ-tubulin
Scale bars 10 µm, t = 30 min.

**Supplementary Figure 3.** XMAP215 and γ-tubulin in combination promote microtubule bundling.

Historically, the mechanism of spontaneous tubulin assembly has been explored using the turbidity data across a range of initial tubulin concentrations [3]. Flyvbjerg and coworkers normalized light scattering to the maximum value at which the data plateaus and normalized time to the time at which turbidity reaches one-tenth its maximum. Finding that the curves were identical following these normalizations, they determined that tubulin follows a single mechanism during assembly, regardless of initial concentration.

We applied this type of analysis to explore the mechanism of assembly in the presence of γ-tubulin and XMAP215. When γ-tubulin is added at 300 nM in combination with XMAP215 at sufficiently high concentration (~40 nM), light scattering increases indefinitely. Normalization of the light scattering of this condition results in a curve with differing shape, specifically with a distinct linear increase in light scattering after turbidity reaches 60% of its maximum value, after approximately 2 min (Supplementary Figure 3A).

To determine if the species of microtubules were different when both γ-tubulin and XMAP215 were present, we performed microtubule pelleting at various time points. By the earliest time point of 3 min, small (≤ 10 microns) microtubule bundles were rare but visible (Supplementary Figure 3B). By the latest time point of 30 min, most microtubules were present in bundles (Supplementary Figure 3C).

The combination of γ-tubulin and XMAP215 in purified solutions promotes bundling of existing microtubules, as noted previously [25]. Comparisons for the effect on nucleation are taken when turbidity reaches 10% of its maximum value, well before bundling occurs.
